# Gyral crowns contribute to the cortical infrastructure of human face processing

**DOI:** 10.1101/2025.03.20.644439

**Authors:** Ethan H. Willbrand, Joseph P. Kelly, Xiayu Chen, Zonglei Zhen, Guo Jiahui, Brad Duchaine, Kevin S. Weiner

## Abstract

Neuroanatomical features across spatial scales contribute to functional specialization and individual differences in behavior across species. Among species with gyrencephalic brains, gyral crown height, which measures a key aspect of the morphology of cortical folding, may represent an anatomical characteristic that importantly shapes neural function. Nevertheless, little is known about the relationship between functional selectivity and gyral crowns—especially in clinical populations. Here, we investigated this relationship and found that the size and gyral crown height of the middle, but not posterior, face-selective region on the fusiform gyrus (FG) was smaller in individuals with developmental prosopagnosia (DPs; *N* = 22, 68% female, aged 25-62) compared to neurotypical controls (NTs; *N =* 25, 60% females, aged 21-55), and this difference was related to face perception. Additional analyses replicated the relationship between gyral crowns and face selectivity in 1,053 NTs (55% females, aged 22-36). These results inform theoretical models of face processing while also providing a novel neuroanatomical feature contributing to the cortical infrastructure supporting face processing.

**Significance Statement:** Understanding how brain structure supports specialized brain functions is a central goal of neuroscience. Here, we identified a role of gyral crown height—an understudied cortical feature—in shaping the cortical infrastructure underlying face processing. By examining face-selective regions of the fusiform gyrus in both neurotypical individuals and those with developmental prosopagnosia, we demonstrate that reduced gyral crown height is associated with diminished face-selective region surface area and impaired face recognition ability. Furthermore, this structural-functional relationship extends to a large neurotypical sample of over 1,000 individuals, highlighting a generalizable link between cortical anatomy and functional specialization. These findings introduce a new neuroanatomical factor to theoretical models of face perception, which could extend to additional neurodevelopmental disorders and other cognitive tasks.

## Introduction

Charting the relationship among brain structure, brain function, and behavior, as well as how these relationships differ in neurodevelopmental disorders, are primary goals in neuroscience. Developmental prosopagnosia (DP) is a particularly intriguing neurodevelopmental disorder to investigate these relationships as individuals with DP have profound deficits in face recognition despite normal low-level vision, normal intelligence, and no explicit insult to the brain (Susilo and Duchaine, 2013; Behrmann et al., 2016). Broadly speaking, the fusiform face area (FFA) (Kanwisher et al., 1997; Kanwisher, 2010), which is a widely studied functional region on the lateral fusiform gyrus (FG) that is causally involved in face perception (Parvizi et al., 2012; Rangarajan et al., 2014; Duchaine and Yovel, 2015; Schalk et al., 2017; Jonas et al., 2018; Jonas and Rossion, 2021), is often a functional neuroanatomical target in DP studies.

However, ongoing research shows that there are at least two functionally and structurally distinct face-selective regions on the FG in neurotypical controls (NTs) (Pinsk et al., 2009; Weiner and Grill-Spector, 2010; Weiner et al., 2010, 2014, 2016, 2017a; Davidenko et al., 2012; Julian et al., 2012; Kietzmann et al., 2012; Parvizi et al., 2012; Çukur et al., 2013; Engell and McCarthy, 2013; McGugin et al., 2014, 2015, 2016; Gomez et al., 2015, 2017, 2018; Kay et al., 2015; Stigliani et al., 2015, 2019; Zhen et al., 2015; Natu et al., 2016, 2019; Elbich and Scherf, 2017; Scherf et al., 2017; Rosenke et al., 2020, 2021; Finzi et al., 2021; Nordt et al., 2021; Chen et al., 2023; Weiner and Willbrand, 2023). Indeed, one such study recently showed that two FG face-selective regions—pFus-faces/FFA-1 and mFus-faces/FFA-2—were structurally and functionally distinct based on architectural, functional, and connectivity features in over 1,000 participants (Chen et al., 2023). Despite these dozens of studies in NTs, to our knowledge, no study has tested the structural, functional, and behavioral relevance of these distinct FG face-selective regions in DPs and NTs.

To fill this gap in knowledge, we first manually identified FG face-selective regions in 47 participants [22 DPs (15 females, ages 25-62) and 25 NTs (15 females, ages 21-55)] to determine whether the incidence rates of FG face-selective regions differed between NTs and DPs. Second, using prior criteria (Chen et al., 2023), we assessed whether the topographical patterning of the FG face-selective regions differed between DPs and NTs. Third, we extracted neuroanatomical and morphological features (gyral height, cortical thickness, surface area) of each region and tested for group-related differences. We targeted gyral height as recent research showed a relationship between genetics and cortical curvature (gyral crowns in face-selective regions and sulcal pits in place-selective regions) in ventral temporal cortex (Abbasi et al., 2020). Fourth, we tested whether neuroanatomical properties correlated with the size of FG face-selective regions. Fifth, we examined whether the extracted properties of the FG face-selective regions were related to individual differences in face perception ability (Duchaine and Nakayama, 2006), and if neuroanatomical properties mediated the relationship between face-selectivity and behavior. Sixth, we tested whether any structure-function relationships identified in this sample extended to a separate, larger sample from the Human Connectome Project (*N*=1053; (Chen et al., 2023).

This multi-pronged approach showed that (i) the incidence of FG face-selective regions does not differ between DPs and NTs, (ii) the gyral crown of both FG face-selective regions, but not cortical thickness, differed between groups, (iii) the size of mFus-faces/FFA-2, but not pFus-faces/FFA-1, differs between DPs and NTs, (iv) there was a relationship between gyral crowns and FG face-selective regions in DPs, NTs, and the larger HCP sample, (v) the size of mFus-faces/FFA-2, but not pFus-faces/FFA-1 is related to face perception ability, and (vi) the latter relationship was mediated by the gyral crown of mFus-faces/FFA-2. These results inform theoretical models of face processing while also providing a novel neuroanatomical feature (gyral crowns) contributing to the cortical infrastructure supporting face processing.

## Materials and Methods

### Experimental Model and Subject Details

#### Participants

##### Dataset 1

Twenty-two DPs (seven males, mean age = 41.9 years old) and 25 NTs (10 males, mean age = 42.3 years old) participated in the study. DPs were recruited from www.faceblind.org, and all reported problems in daily life with face recognition. To assess their face recognition, DPs were tested with the Cambridge Face Memory Test (CFMT) (Duchaine and Nakayama, 2006), a famous face test (Duchaine and Nakayama, 2005), and an old–new face discrimination test (Duchaine and Nakayama, 2005). All but one DP performed two or more standard deviations (SDs) below the mean of published control results in at least two of the three diagnostic tests (Duchaine et al., 2007a, 2007b). The DP participant who did not reach −2 SDs on two tests scored poorly on two of the three tasks (CFMT: z = −1.9; famous face: z = −7.1; old– new: z = −0.5), so we included them to increase the sample size. All participants had normal or corrected-to-normal vision and had no current psychiatric disorders. Participants provided written informed consent before doing the tasks, and all procedures were approved by Dartmouth’s Committee for the Protection of Human Participants. Participants were leveraged from a prior study on category selectivity in DP (Jiahui et al., 2018) and on ventral temporal sulcal morphology in DP (Parker et al., 2023).

##### Dataset 2

Data for the Human Connectome Project neurotypical adult cohort (HCP) analyzed in the present study were sourced from the freely available HCP database (https://humanconnectome.org/study/hcp-young-adult) (Van Essen et al., 2012). The dataset consisted of 1053 participants (575 females, ages 22 to 36). These data were previously acquired using protocols approved by the Washington University Institutional Review Board.

## Method Details

### Face recognition task

#### Dataset 1

In the present study, we focused on the CFMT as our quantitative measure of face recognition for three main reasons. First, the CFMT is a commonly used measure of unfamiliar face recognition (Duchaine and Nakayama, 2006). Second, the CFMT is a well-established test with high reliability: previous studies show a Spearman-Brown split-half reliability for CFMT of .91, as well as a test-retest reliability of .70 and alternate forms of reliability of .76 (Wilmer et al., 2010, 2012). Third, we focus on the CFMT to maximize the number of participants included in the present measurements. Specifically, the majority of participants (except one NT) completed the CFMT.

### MRI data

#### Brain data acquisition

*Dataset 1:* All participants were scanned in a 3.0T Philips MRI scanner (Philips Medical Systems, WA, USA) with a SENSE (SENSitivity Encoding) 32-channel head coil. A high-resolution anatomical volume was acquired at the beginning of the scan using a high-resolution 3D magnetization-prepared rapid gradient-echo sequence (220 slices, field of view = 240 mm, acquisition matrix = 256 x 256, voxel size = 1 x 0.94 x 0.94 mm).

*Dataset 2:* Anatomical T1-weighted MRI scans (0.7-mm voxel resolution) were obtained in native space from the HCP database, along with outputs from the HCP-modified FreeSurfer pipeline (v5.3.0) (Dale et al., 1999; Fischl et al., 1999a; Glasser et al., 2013). Additional details on image acquisition parameters and image processing can be found in (Glasser et al., 2013). All subsequent sulcal labeling and extraction of anatomical metrics were calculated from the cortical surface reconstructions of individual participants generated through the HCP’s custom modified version of the FreeSurfer pipeline (Glasser et al., 2013).

#### Cortical surface reconstruction

Cortical surface reconstructions were generated for each participant from their T1 scans using the standard FreeSurfer pipeline (v6.0.0; https://surfer.nmr.mgh.harvard.edu) (Dale et al., 1999; Fischl et al., 1999a, 1999b). Cortical reconstructions were created from the resulting boundary made by segmenting the gray and white matter in each anatomical volume with FreeSurfer’s automated segmentation tools (Dale et al., 1999). Subsequent ROI labeling and extraction of morphological metrics were calculated from individual participants’ cortical surface reconstructions. This process was carried out blind to the participant group.

#### Functional localizer

*Dataset 1:* A detailed description of the functional scanning parameters has been previously described in (Jiahui et al., 2018). Here, we provide a brief overview. Participants completed a one-back task during a dynamic localizer scan containing five visual categories (faces, scenes, bodies, objects, and scrambled objects). Each participant completed five scans, composed of 10 12-s category blocks of video clips interleaved with 12-s fixation blocks (4.2 minutes in total). Each visual category was displayed twice in each scan in a quasi-random order. In each category block, six 1,500-ms video clips were presented interleaved by blank fixation screens presented for 500 ms. Stimuli were presented using SuperLab 4.5.3 (https://cedrus.com/superlab/index.htm) and displayed to the participant via a Panasonic DT-4000UDLP projector (resolution: 1,024 × 768; refresh rate: 60 Hz) at the rear of the scanner.

The five runs were then divided into localization runs and test runs to carry out a “leave-one-out” analysis (Norman-Haignere et al., 2013, 2016). In each of the leave-one-out combinations, four of the five runs for a participant were used to localize the vertices that showed the strongest preference for the preferred category. To avoid “double-dipping” (Kriegeskorte et al., 2009), the responses of the selected voxels to each stimulus condition were then measured in the left-out run. All five combinations were analyzed and then averaged to produce the final result for each participant. Finally, face-selective maps were created from the difference between the response to faces and the response to objects for each participant.

*Dataset 2:* Face-selective regions in HCP participants were localized using a working memory task in which four stimulus types (faces, places, tools, and body parts) were presented in separate blocks (Barch et al., 2013). The localizer consisted of two runs, and each run contained eight task blocks (10 trials of 2.5 s each, for 25 s) and 4 fixation blocks (15 s each). Within each run, half of the task blocks used a 2-back working memory task and the other half implemented a 0-back working memory task. A 2.5 s cue indicated the task type at the start of the block. For each trial, the stimulus was presented for 2 s, followed by a 500 ms inter-trial interval (ITI). Linear contrasts were computed to estimate effects of interest (e.g., faces vs. other categories). Fixed-effects analyses were conducted to estimate the average effects across runs within each participant.

### Manual definition of face-selective regions

Face-selective regions on the lateral fusiform gyrus (FG) were manually delineated for each hemisphere and each participant based on individual, thresholded (top 5% of face-selective vertices) face-selective activation maps (faces versus objects) created for each participant (**Fig. 1A**) (Jiahui et al., 2018). From this thresholded map, regions of interest (ROIs) were labeled as either mFus-faces/FFA-2 or pFus-faces/FFA-1 based on previously published criteria differentiating the cortical location of the two regions relative to sulci within and surrounding the FG (**Fig. 1A**) (Weiner et al., 2014; Weiner, 2019; Chen et al., 2023). Specifically, mFus-faces/FFA-2 is located adjacent to the anterior tip of the mid-fusiform sulcus (MFS), whereas pFus-faces/FFA-1 is located on the posterior aspect of the FG, extending into the occipito-temporal sulcus (**Fig. 1A**) (Weiner et al., 2014; Weiner, 2019; Chen et al., 2023). To define each region in the DP and NT sample, K.S.W. identified each region manually on the individual thresholded face-selective map and then authors E.H.W. and J.P.K. labeled these regions in FreeSurfer using *tksurfer* tools (https://surfer.nmr.mgh.harvard.edu/fswiki/TkSurfer). As in prior work (Chen et al., 2023), we used a liberal threshold (top 5% of face-selective vertices) for the main reason that we did not want to artificially inflate the “separate” group (described below) by using a strict threshold (e.g., top 2.5%) or the “continuous” group (described below) by using a too liberal threshold (e.g., top 10%). Crucially, these regions were defined blind to the identity of all participants (i.e., whether or not they were an NT or DP). The inflated cortical surfaces presented in **Figure 1** were generated with Freeview (https://surfer.nmr.mgh.harvard.edu/fswiki/FreeviewGuide). The FG face-selective regions were defined in the HCP sample in our prior work by X.C. and K.S.W. (Chen et al., 2023).

**Figure 1.**
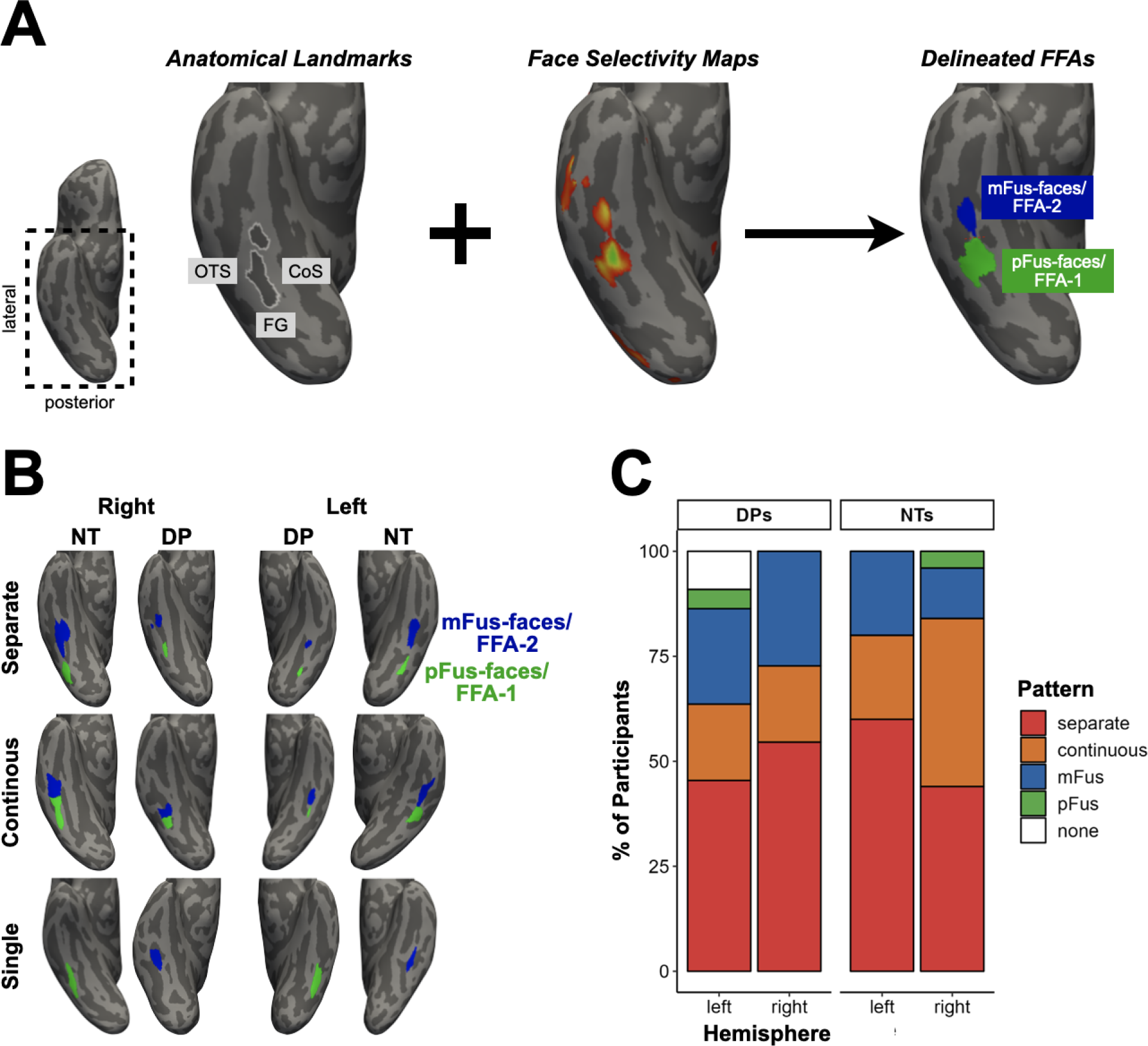
Topographical patterning of pFus-faces/FFA-1 and mFus-faces/FFA-2 does not differ between DPs and NTs. **A.** Methodology to identify pFus-faces/FFA-1 and/or mFus-faces/FFA-2 in individual participants. Inflated cortical surface reconstruction of a right hemisphere (inset for the location of zoomed portion) from an example NT participant displaying the process for identifying face-selective regions on the FG at the individual level. Face-selective regions were manually defined on the lateral FG in 22 DPs and 25 NTs by using individual anatomical landmarks [OTS: occipitotemporal sulcus; CoS: collateral sulcus; MFS: mid-fusiform sulcus (white outline)] and face-selective maps (middle hemisphere; faces versus objects, top 5% of vertices). These FG face-selective regions were labeled as either mFus-faces/FFA-2 or pFus-faces/FFA-1 in each hemisphere based on previously published criteria (Weiner et al., 2014). **B.** FG face-selective regions are displayed for 12 randomly selected hemispheres (one example of each pattern in each hemisphere for each group). Hemispheres are zoomed in on the ventral temporal cortex as in A. Top row: separate group; Middle row: continuous group; Bottom row: single group. Left two columns: right hemispheres; Right two columns: left hemispheres. For each pattern and hemisphere, the outer hemisphere is from an NT, whereas the inner hemisphere is from a DP. **C.** Bar plot visualizing the incidence of the topographical patterns (colors; see key) as a function of hemisphere (x-axis) and group (left: DPs; right: NTs). The incidence (0, 1, or 2 regions) and pattern (see key) of the face-selective regions did not significantly differ between groups (all p-values > .05).

### Quantifying the incidence rates and pattern

We categorized the spatial organization of mFus-faces/FFA-2 and pFus-faces/FFA-1 into three types, or topological groups as in prior work in over 1,000 participants (Chen et al., 2023): separate, continuous, and single. The “separate” group consisted of two cortically distinct face-selective regions in a given hemisphere that were separated by a cortical gap (**Fig. 1B**). The “continuous” group consisted of two regions that were identifiable and contiguous, but could be separated based on previously proposed anatomical criteria using cortical folding (**Fig. 1B**) (Weiner et al., 2014; Weiner, 2019; Chen et al., 2023). Note that the most likely factor contributing to the two regions being continuous is the spatial coarseness of the BOLD signal.

That is, there is likely a cortical gap in these individuals, but the coarseness of the spatial spread of the BOLD signal causes the two regions to blur together (Chen et al., 2023). The “single” group consisted of one region in which either mFus-faces/FFA-2 or pFus-faces/FFA-1, but not both, was identifiable in a given hemisphere (**Fig. 1B**). After determining these three groups, we summarized the incidence rate of each group (DP, NT) by quantifying how many hemispheres had (i) two or < 2 face-selective regions and (ii) each topological type.

### Quantifying surface area, thickness, and gyral crown height

As in prior work (Chen et al., 2023), the surface area (in squared millimeters) and average cortical thickness (in millimeters) of each region were extracted with the mris_anatomical_stats function in FreeSurfer (Fischl and Dale, 2000). The maximum gyral height (“gyral crown”; quantified as the minimum value of the FreeSurfer .sulc file for each FG face-selective region; (Dale et al., 1999; Fischl et al., 1999a, 1999b) of each region was extracted using custom Python code leveraging various functions from the numpy, pandas, nilearn, and nibabel packages (from (Miller et al., 2021). Note that values in the .sulc file are calculated based on how far removed a vertex is from what is referred to as a “mid-surface,” which is determined computationally so that the mean of the displacements around this “mid-surface” is zero. Thus, generally, gyri have negative values, while sulci have positive values. Therefore, the corresponding maximum gyral height of each region was quantified based on this assignment.

### Quantification and Statistical Analysis

All statistical tests and data visualization were implemented in R (v4.0.1; https://www.r-project.org/) via RStudio (RStudio v.2021.09.0+351; https://posit.co/). Fisher’s exact tests were carried out to compare the incidence rates of face-selective regions and the three topographical types between groups with the fisher.test function from the stats package. 2-way mixed-model ANOVAs were implemented for each quantitative metric (surface area, gyral crown height, average cortical thickness) with the lme and anova functions from nlme and stats packages for each region (mFus-faces/FFA-2, pFus-faces/FFA-1) separately. Hemisphere (left, right) was a within-participant factor, and group (DP, NT) was a between-participant factor. Effect sizes for the significant effects are reported with the partial eta-squared (η2) metric, computed with the eta_squared function from the effectsize package. Next, we correlated the properties of mFus-faces/FFA-2 and pFus-faces/FFA-1 to face perception ability (as measured by performance on the CFMT; (Duchaine and Nakayama, 2006). Spearman rank-order correlations (r_s_) and p-value calculations were implemented with the cor.test function from the stats package. We used r_s_ to be less influenced by any potential outliers. For each correlation, we first tested for a relationship across hemispheres, followed up by testing within each hemisphere separately. To quantify whether there was an indirect effect of gyral crown height on the relationship between mFus-faces/FFA-2 properties and CFMT scores, we implemented a bootstrapped (with 1,000 simulations) causal mediation analysis with the mediate function from the mediation package. Results for the mediation analyses are reported with the bootstrapped average causal mediation effect (ACME) alongside the 95% confidence interval. Plots were created with the ggplot2 package.

### Data and code availability

Processed data used for the present project have been deposited on GitHub and are publicly available as of the date of publication (https://github.com/cnl-berkeley/stable_projects). All original code used for the present project has also been deposited on Github and is publicly available as of the date of publication. Any additional information required to reanalyze the data reported in this paper is available from the lead contact, Kevin Weiner, upon request.

## Results

### Incidence rates of FG face-selective regions do not differ between DPs and NTs

Manually delineating FG face-selective regions in 22 DPs (44 hemispheres) and 25 NTs (50 hemispheres; **Materials and Methods** and **Fig. 1A, B**) revealed that at least one face-selective region was identifiable in every hemisphere in NTs and in all but two DP hemispheres (**Fig. 1C**). 82% of NTs (Left: 80%; Right: 84%) and 68.18% of DPs (Left: 63.64%; Right: 72.73%; **Fig. 1C**) had two face-selective regions on the FG. As in prior work (Chen et al., 2023), these incidence rates could be further categorized into one of three different types in a given hemisphere: separate, continuous, or single (**Materials and Methods**; **Fig. 1B**). The most common was the separate group, in which 52% of NT hemispheres (LH: 60%; RH: 44%) and 50% of DP hemispheres (LH: 45.46%; RH: 54.55%) contained two face-selective regions that were separated by a cortical gap of several millimeters (**Fig. 1B, C**). In the continuous group, which consisted of 30% of cases in NTs (LH: 20%; RH: 40%) and 18.18% of cases in DPs (LH: 18.18%; RH: 18.18%), mFus-faces/FFA-2 and pFus-faces/FFA-1 were identifiable and largely contiguous (**Fig. 1B, C**). Specifically, the former was identified as the functional region located adjacent to the anterior tip of the mid-fusiform sulcus (MFS), while the latter was identified as the functional region located adjacent to the posterior extent of the MFS extending into the lateral FG and the nearby occipitotemporal sulcus (**Fig. 1**) (Weiner et al., 2014; Weiner, 2019; Chen et al., 2023). In 18% of NT cases (LH: 20%; RH: 16%) and 27.27% of DP cases (LH: 27.27%; RH: 27.27%), either mFus-faces/FFA-2 or pFus-faces/FFA-1, but not both, was identifiable in a given hemisphere based on the criteria just described (**Fig. 1B, C**). A single mFus-faces-FFA/2 was more common than a single pFus-faces/FFA-1 in both NTs (*mFus*: LH: 20%; RH: 12%; *pFus*: LH: 0%; RH: 4%; Χ^2^ = 5.44, *p* = 0.01) and DPs (*mFus*: LH: 22.73%; RH: 27.27%; *pFus*: LH: 4.55%; RH: 0%; Χ^2^ = 8.33, *p* = .003). In the left hemisphere, two DPs (9.09%) did not have an identifiable mFus-faces/FFA-2 or pFus-faces/FFA-2 (**Fig. 1C**). These results are similar to a recent study from our laboratory (Chen et al., 2023), which found that the highest percentage of participants was in the separate group, then the continuous group, and then the single group. The percentages of mFus-faces/FFA-2 and pFus-faces/FFA-1 combinations did not differ between DPs and NTs (Both: p = .21; Left: p = .50; Right: p = .26; **Fig. 1C**).

### Gyral crown and surface area of FG face-selective regions differs between DPs and NTs

As recent findings show a relationship between cortical folding in relation to category-selective regions in high-level visual cortex (Weiner et al., 2014; Abbasi et al., 2020; Arcaro et al., 2020; Natu et al., 2021), we tested if the gyral crown (highest point of cortical folding) of each FG face-selective region differed between groups. We also considered cortical thickness as it is a common measure examined in FG face-selective regions (Bi et al., 2014; McGugin et al., 2016, 2020; Zebrowitz et al., 2016; Chen et al., 2023). Crucially, these features were selected based on the functional boundaries of each region, and thus, are not necessarily dependent on any given cortical landmark (e.g., a specific sulcus or gyrus). Mixed-model ANOVAs for each region (mFus-faces/FFA-2, pFus-faces/FFA-1) with hemisphere (left, right), and group (NT, DP) as factors revealed a main effect of group on gyral crown height for each region (mFus-faces/FFA-2: F(1, 45) = 10.68, p = .002, η2 = 0.19; pFus-faces/FFA-1: F(1, 42) = 11.90, p = .001, η2 = 0.22), such that the gyral crown height of both FG face-selective regions was lower in DPs (mFus-faces/FFA-2: mean ± se = −3.49 ± 0.36; pFus-faces/FFA-1: mean ± se = −4.07 ± 0.41) compared to NTs (mFus-faces/FFA-2: mean ± se = −5.02 ± 0.27; pFus-faces/FFA-1: mean ± se = −6.38 ± 0.36; **Fig. 2A**). There were no main effects of hemisphere or hemisphere and group interactions on gyral crown height for either region (ps > .28; **Fig. 2A**). In contrast to the main effect of gyral crown height, there was no main effect of group on cortical thickness for either region (ps > .36; **Extended Data Fig. 1A**). There were also no main effects of hemisphere or hemisphere and group interactions on cortical thickness for either region (ps > .39; **Extended Data Fig. 1A**).

**Figure 2.**
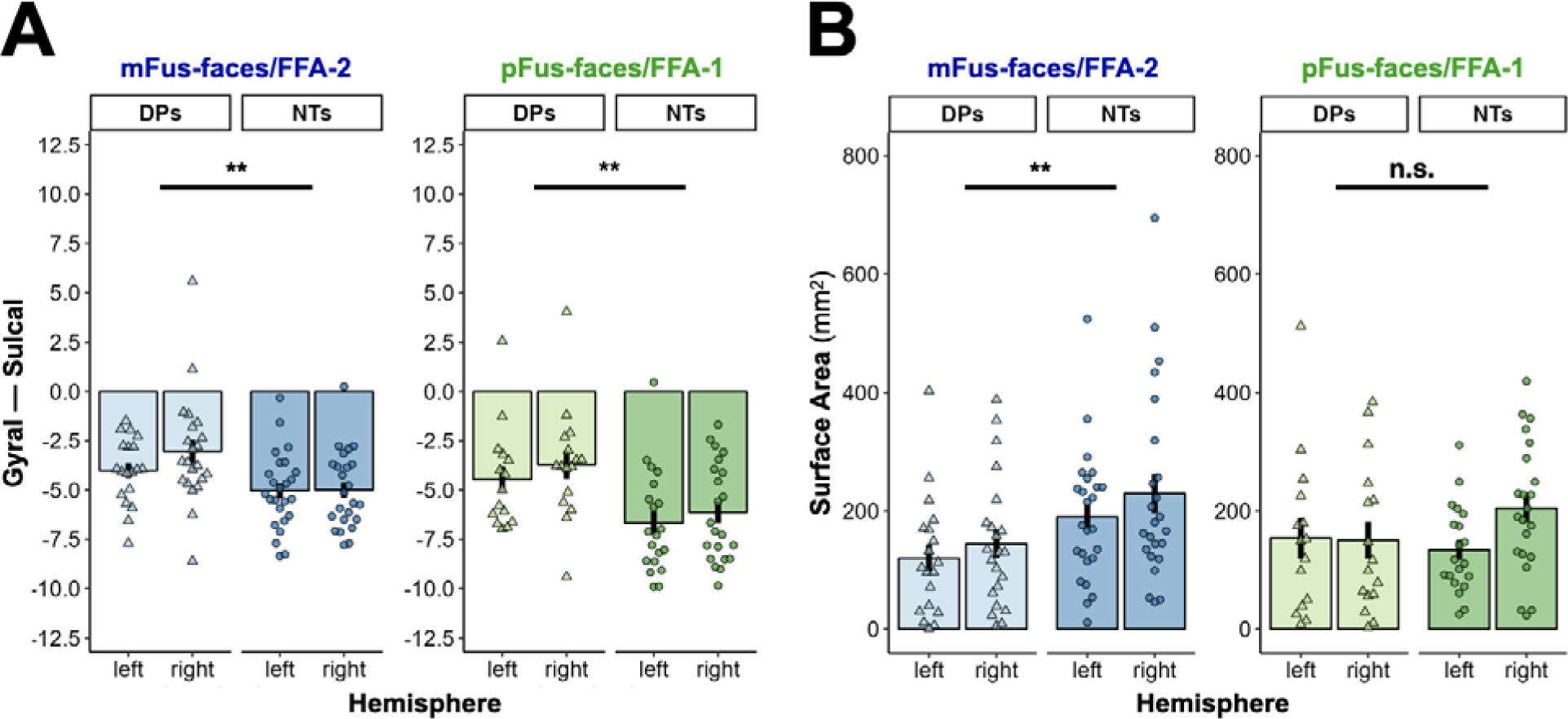
Gyral crown height and surface area of FG face-selective regions differ between DPs and NTs. **A.** Bar plots and error bars (mean ± se) visualizing region maximal gyral height (“gyral crown”; in FreeSurfer units: the more negative value indicates higher gyral height) as a function of hemisphere (x-axis) and group (left: DPs; right: NTs) for both mFus-faces/FFA-2 (left plot) and pFus-faces/FFA-1 (right plot). Individual dots represent individual values for each participant (DPs = triangles, NTs = circles). The horizontal line and asterisks visualize the statistical significance of the group difference for gyral crown across groups. **B.** Same as A, but for region surface area (in mm^2^). (**, p < .01; n.s., p > .05).

We also wanted to examine whether the morphological properties (i.e., surface area) of the FG face-selective regions differed between groups. To do so, we next implemented mixed-model ANOVAs for each region with factors of hemisphere (left, right) and group (NT, DP) to test whether the size (surface area in mm^2^) of each FG face-selective region differed between groups. The two DP hemispheres without an FG face-selective region were not included in analyses. These analyses revealed two main findings. First, there was a main effect of group on the size of mFus-faces/FFA-2 (F(1, 45) = 8.00, p = .007, η2 = 0.15), such that the surface area was generally smaller in DPs (mean ± se = 133 ± 15.9 mm^2^) compared to NTs (mean ± se = 209 ± 19.5 mm^2^; **Fig. 2B**, left). Second, there was no main effect of group on the size of pFus-faces/FFA-1 (F(1, 42) = 0.53, p = .47, η2 = 0.01; DP: mean ± se = 152 ± 19.2 mm^2^; NT: mean ± se = 171 ± 14.2 mm^2^;**Fig. 2B**, right). This difference in size is consistent with a previous preprint in a smaller group of DPs (N = 7) showing that the volume of mFus-faces/FFA-2, but not pFus-faces/FFA-1, was smaller in DPs compared to NTs (Witthoft et al., 2016). There was also no main effect of hemisphere or hemisphere and group interaction for either region (ps > .14; **Fig. 2B**).

### Gyral crown height correlates with the surface area of FG face-selective regions

Next, we tested if the neuroanatomical and morphological features of FG face-selective measures were correlated with the surface area of FG face-selective regions. Gyral crown height was related to the size of both regions across hemispheres, such that the higher the gyral crown within the anatomical locus within the FG, the larger the surface area of the face-selective region (mFus-faces/FFA-2: r_s_ = −0.44, p = .00001; pFus-faces/FFA-1: r_s_ = −0.30, p = .01; **Fig. 3A**). In both regions, this relationship was stronger in the right hemisphere (mFus-faces/FFA-2: r_s_ = −0.59, p = .0001; pFus-faces/FFA-1: r_s_ = −0.36, p = .025) compared to the left (mFus-faces/FFA-2: r_s_ = −0.29, p = .053; pFus-faces/FFA-1: r_s_ = −0.26, p = .13; **Fig. 3A**). Conversely, cortical thickness was unrelated to the size of either region (ps > .47; **Extended Data Fig. 1B**).

**Figure 3.**
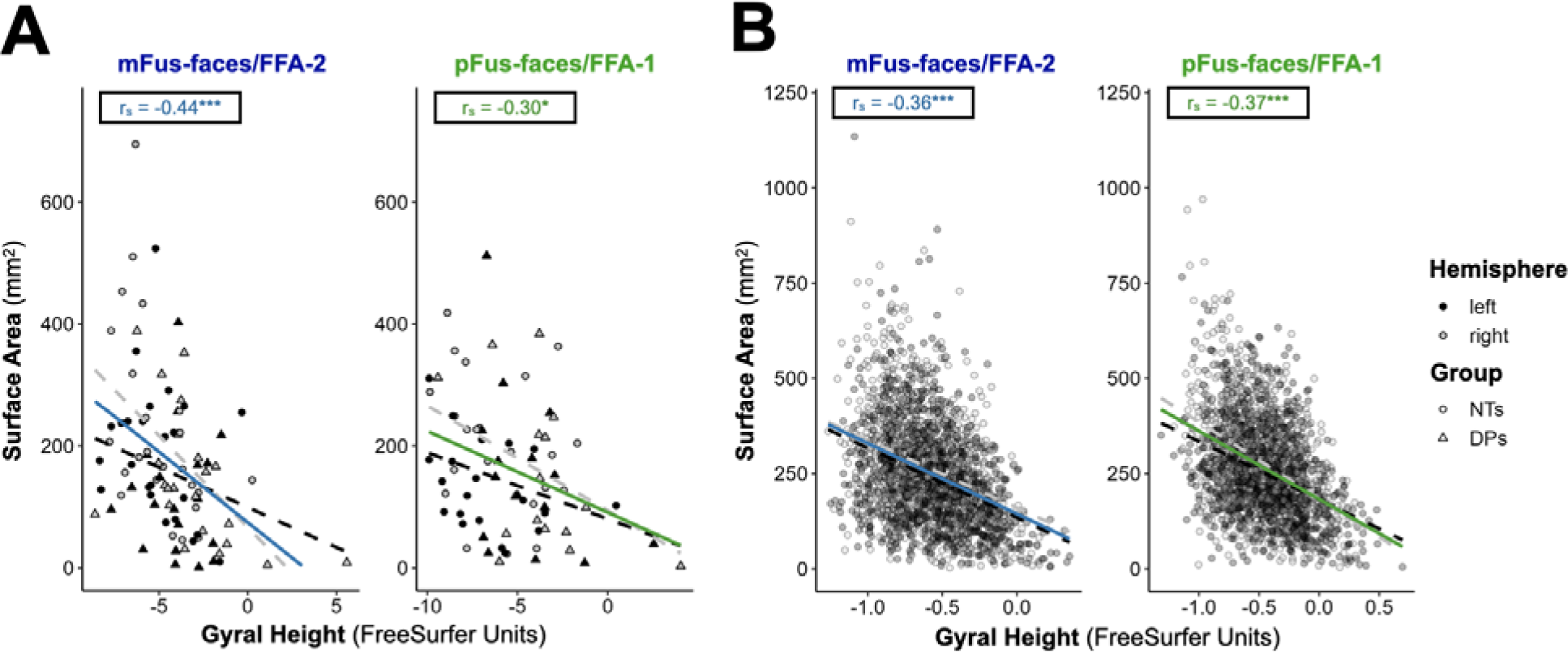
Gyral crown height is related to the surface area of FG face-selective regions. **A.** Scatterplot of maximal gyral height (“gyral crown”; in FreeSurfer units: the more negative value indicate higher gyral height; x-axis) and surface area (in mm^2^; y-axis) for mFus-faces/FFA-2 (left) and pFus-faces/FFA-1 (right). The solid, colored line corresponds to the best-fit line across hemispheres alongside the corresponding correlation coefficient (r_s_) and p*-*values (asterisks). The best-fit lines for the left and right hemispheres are also shown as dashed lines colored according to the key. Individual dots represent individual values for each participant which are colored by hemisphere (left = black, right = white; see key) and shaped according to the group of that individual (DPs = triangles, NTs = circles; see key). **B.** Same format as A, but for the HCP sample (N = 1053) (Chen et al., 2023). (***, p < .001; **, p < .01; * p < .05).

To test whether this novel relationship between gyral crowns and the surface area of FG face-selective regions was also present in a larger sample that used a different methodology (e.g., task details, statistical threshold, etc.), we leveraged the prior definitions of FG face-selective regions from 1053 participants (478 males, mean age = 29 years old; 2106 hemispheres) from the Human Connectome Project (HCP; for additional sample and methodological details see (Chen et al., 2023). This analysis showed that the relationship between gyral crown height and the size of FG face-selective regions was replicated for both mFus-faces/FFA-2 (across hemispheres: r_s_ = −0.36, p < 2.2e-16; left hemisphere: r_s_ = −0.38, p < 2.2e-16; right hemisphere: r_s_ = −0.33, p < 2.2e-16; **Fig. 3B**, left) and pFus-faces/FFA-1 (across hemispheres: r_s_ = −0.37, p < 2.2e-16; left hemisphere: r_s_ = −0.36, p < 2.2e-16; right hemisphere: r_s_ = −0.39, p < 2.2e-16; **Fig. 3B**, right). The relationship between cortical thickness and size was weak for both regions in the HCP sample (**Extended Data Fig. 1C**).

### The size of mFus-faces/FFA-2, but not pFus-faces/FFA-1, correlates with face processing ability and is mediated by gyral crown height

To build upon prior work showing a relationship between face processing ability and the size of the FFA (Golarai et al., 2007, 2010; Furl et al., 2011; Elbich and Scherf, 2017), but which did not consider individual FG face-selective regions, we tested whether the size of each FG face-selective region was also correlated with performance on the Cambridge Face Memory Test (CFMT) (Duchaine and Nakayama, 2006). In addition, if a correlation was significant, we assessed if this relationship was mediated by gyral crown height. Importantly, our voxel selection criteria for ROI definitions were defined a priori and were completely independent of the behavioral measures. Concomitantly, the relationship between the surface area of face-selective regions and CFMT performance is orthogonal to the voxel-selection process (Nichols and Poline, 2009; Poldrack and Mumford, 2009; Vul et al., 2009).

For mFus-faces/FFA-2, there was a positive relationship between CFMT performance and ROI size, such that the larger the surface area, the better face processing performance (across hemispheres: r_s_ = 0.33, p = .001; left hemisphere: r_s_ = 0.38, p = .011; right hemisphere: r_s_ = 0.31, p = .041; **Fig. 4A**, left). By contrast, there was not a significant relationship between CFMT performance and the size of pFus-faces/FFA-1 (across hemispheres: r_s_ = 0.13, p = .25; left hemisphere: r_s_ = 0.008, p = 0.96; right hemisphere: r_s_ = 0.25, p = .13; **Fig. 4A**, right). The size correlations were modestly different between regions (across hemispheres: Fisher’s Z = 1.31, p = .095; left hemisphere: Fisher’s Z = 1.66, p = .049; right hemisphere: Fisher’s Z = 0.28, p = .38). Interestingly, these findings are similar to a previously published pre-print in a much smaller sample size of DPs (N = 7) in which there was a marginally significant relationship between the volume of mFus-faces/FFA-2 and CFMT (r = .49, p = .08), but not between the volume of pFus-faces/FFA-1 and CFMT (r = .28, p = .32) (Witthoft et al., 2016).

**Figure 4.**
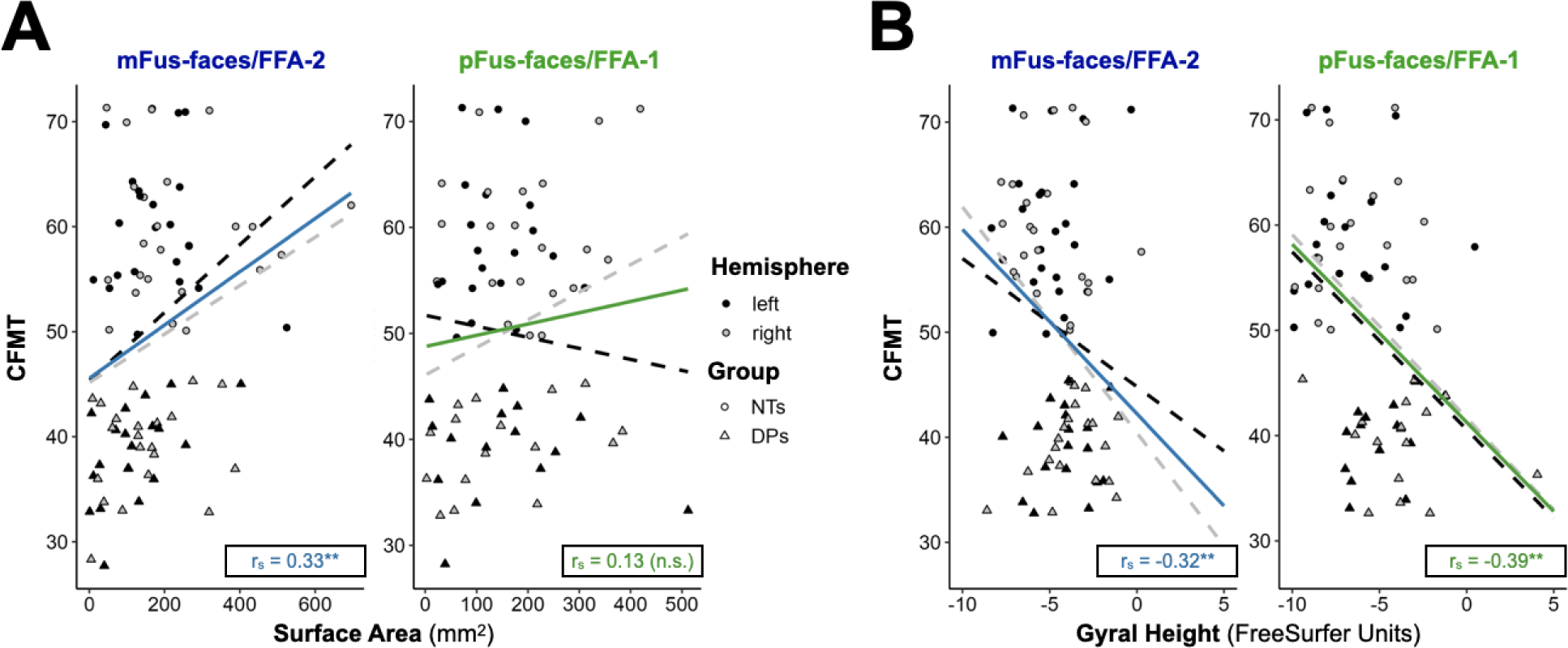
Size and gyral crown height of FG face-selective regions selectively correlate with face recognition ability. **A.** Scatterplot of surface area (in mm^2^; x-axis) and Cambridge Face Memory Test (CFMT; y-axis) performance for mFus-faces/FFA-2 (left) and pFus-faces/FFA-1 (right). The solid, colored line corresponds to the best-fit line across hemispheres alongside the corresponding correlation coefficient (r_s_) and p*-*values. The best-fit lines for the left and right hemispheres are also shown as dashed lines colored according to the key. Individual dots represent individual values for each participant which are colored by hemisphere (left = black, right = white; see key) and shaped according to the group of that individual (DPs = triangles, NTs = circles; see key). **B.** Same as A, but for maximal gyral crown height (in FreeSurfer units: the more negative value indicates higher gyral crown height). (**, p < .01; n.s., p > .05).

Examining the relationship between the gyral crown height of mFus-faces/FFA-2 and CFMT performance also revealed a significant overall relationship (across hemispheres: r_s_ = - 0.32, p = .002) with stronger effects in the right hemisphere (r_s_ = −0.39, p = .007) than in the left hemisphere (r_s_ = −0.24, p = 0.11; **Fig. 4B**, left). A similar relationship between the gyral crown height of pFus-faces/FFA-1 and CFMT performance was observed (across hemispheres: r_s_ = - 0.39, p = .007; left hemisphere: r_s_ = −0.40, p = 0.018; right hemisphere: r_s_ = −0.42, p = .009; **Fig. 4B**, right). There was no relationship between cortical thickness of either region and CFMT (ps > .32; **Extended Data Fig. 1D**).

Finally, given the relationship between gyral crown height and the size of mFus-faces/FFA-2 (**Fig. 3**), we sought to determine whether gyral crown height mediated the observed relationship between the size of mFus-faces/FFA-2 and CFMT performance with causal mediation analyses. Computing this indirect effect of gyral crown height from 1,000 bootstrapped simulations with the bias-corrected method revealed that gyral crown height mediated the relationship between the size of mFus-faces/FFA-2 and CFMT (indirect effect [95% CI] = 0.008 [0.0005, 0.02], p = .04).

## Discussion

In the present study, we examined the structure and function of separate FG face-selective regions (mFus-faces/FFA-2 and pFus-faces/FFA-1) in DPs relative to NTs. We showed that the incidence and topographical location of these regions relative to cortical folding was organized similarly between DPs and NTs. The lack of differences regarding the topographical arrangement of the FG face-selective regions is consistent with previous findings showing that DPs generally have a comparable gross functional organization of VTC to NTs (Avidan et al., 2005, 2014; Avidan and Behrmann, 2009; Garrido et al., 2009; Zhang et al., 2015; Jiahui et al., 2018), which we now extend to a finer scale (that is, considering multiple FG face-selective regions and tertiary sulci in VTC).

However, despite these observed topographical similarities, the more anteriorly located mFus-faces/FFA-2 was smaller in DPs compared to NTs, while this was not the case for the more posterior pFus-faces/FFA-1. We further showed that the surface area of mFus-faces/FFA-2, but not pFus-faces/FFA-1, was related to CFMT performance. Our results also indicate a new relationship between gyral crowns and the surface area of FG face-selective regions: gyral crown height, but not cortical thickness, was related to the surface area of FG face-selective regions, and differed between DPs and NTs. This structural-functional effect generalized to a large sample size of 1053 participants from the Human Connectome Project, indicating a new structural-functional relationship of two anatomically and functionally distinct face-selective regions on the FG. Below, we discuss these results in the context of (i) coupling between local anatomical features and category selectivity in ventral temporal cortex, (ii) functional and structural differences between DPs and NTs, and (iii) the role of gyral crown height and face selectivity.

### Coupling between local anatomical features and category selectivity in ventral temporal cortex

The present findings build on recent studies identifying a tight coupling between local anatomical features and category selectivity in medial and lateral aspects of human ventral temporal cortex (VTC). In medial VTC, Natu and colleagues identified a relationship between deep sulcal points (e.g., sulcal pits or sulcal roots) in the collateral sulcus (CoS) and place selectivity in children and young adults (Natu et al., 2021). In lateral VTC, Cachia and colleagues identified a relationship between the location of a region selective for visual words and a gyral gap in the occipitotemporal sulcus (OTS) (Cachia et al., 2018). Building on that work, Bouhali and colleagues showed that longitudinal changes in the OTS gyral gap correlated with longitudinal changes in reading ability—changes that were mediated by underlying white matter properties (Bouhali et al., 2024). Furthermore, previous work also showed a tight coupling between the anterolateral tip of the mid-fusiform sulcus and mFus-faces/FFA-2 (Weiner, 2019). Here, we show that the highest local gyral point within mFus-faces/FFA-2 contributes to this structural-functional coupling between cortical folding and face selectivity.

This coupling between local anatomical features and category selectivity also extends to non-human primates (NHP). For example, there is also a tight coupling between deep sulcal points and place selectivity in VTC (Natu et al., 2021), as well as “bumps” in the superior temporal sulcus and face selectivity in NHPs (Arcaro et al., 2020). Thus, there is growing evidence that local anatomical features are better anatomical predictors of category selectivity in high-level visual cortex across species than gross macroanatomical landmarks (e.g. entire sulci or gyri). Finally, the present findings contribute to work suggesting the importance of gyral crown height and its contributions to brain structure and function (Zhang et al., 2022, 2023), as well as a potential functional relevance of a latitude/longitude coordinate system of sulcal pits and gyral crowns and their relevance to face and place selectivity in human VTC (Auzias et al., 2013), which can also be explored in future work.

### Functional and structural differences between NTs and DPs: Longitudinal white matter tracts?

Our results identify structural and functional differences related to face processing between DPs and NTs focal to mFus-faces/FFA-2 that have cognitive implications. In this section, we consider that differences in underlying white matter properties specific to mFus-faces/FFA-2 could contribute to the differences identified here. For example, there are functionally-defined longitudinal white matter tracts specific to mFus-faces/FFA-2 that are present in both DPs and NTs, but properties of these tracts are different between groups and predictive of face processing ability in NTs, but not DPs (Gomez et al., 2015). Importantly, these tracts are functionally defined and are much smaller than widely studied larger tracts in many neurodevelopmental disorders such as the inferior longitudinal fasciculus. The fact that these tracts are smaller would also support a local difference between groups in which the properties of these tracts specific to mFus-faces/FFA-2 differ between NTs and DPs, but the tracts are still present in each group.

Together, the differences in the underlying white matter of mFus-faces/FFA-2 (Gomez et al., 2015), as well as the functional and anatomical differences identified here, between NTs and DPs could be reflective of a posterior-anterior gradient. For example, Jiahui and colleagues (Jiahui et al., 2018) did not find significant differences in the selectivity of the most posterior face-selective region in VTC on the inferior occipital gyrus (i.e., the occipital face area, OFA). Here, we expand on those findings and show that the largest difference between DPs and NTs is in the more anteriorly positioned, mFus-faces-FFA-2, compared to the more posteriorly positioned pFus-faces-FFA-1. Nevertheless, it is worth noting that other DP studies have identified abnormalities (Zhao et al., 2016, 2018; Tian et al., 2020). As such, the combination of these findings indicate that there may be smaller group differences in posterior face-selective regions compared to anterior face-selective regions in VTC which leads to a targeted hypothesis for future research: the percentage of DPs with abnormalities in VTC face-selective regions will rise as one moves forward in the face processing stream. Thus, an immediate goal of future studies is to further understand why the anterior tip of the MFS extending into the lateral mid-fusiform gyrus has the largest difference between NTs and DPs and if this is a general finding across other populations with difficulties in face processing.

### The role of gyral crown height and face selectivity: Vertical white matter tracts?

Consistent with the previous section relating cortical folding, underlying white matter architecture, and face processing, there is a general anatomical relationship between white matter architecture and cortical folding. For example, long-range white matter fibers have a “gyral bias” (also referred to as “gyral blades”) (Van Essen et al., 2014; Cottaar et al., 2021). An intriguing possibility is that the relationship among gyral crown height, functional features of mFus-faces/FFA-2, and face processing ability indicates that this anatomical location is an anatomical locus that integrates information across cortical networks. For example, recent electrical brain stimulation (EBS) studies support this possibility: EBS delivered to the more posterior pFus-faces/FFA-1 shows effects that are specific to perceptual aspects of faces, while EBS delivered to the more anterior mFus-faces/FFA-2 shows effects supporting an integration between networks that are related to semantics such as naming and person knowledge (Parvizi et al., 2012; Rangarajan et al., 2014; Schrouff et al., 2020).

In addition to the long-range longitudinal white matter tracts discussed in the previous section, there are two major long-range vertical white matter tracts that could also contribute to this functional dissociation between pFus-faces/FFA-1 and mFus-faces/FFA-2: the vertical occipital fasciculus (VOF) and the arcuate fasciculus, which consists of separate vertical and arching components (Yeatman et al., 2014; Weiner et al., 2017b). The VOF terminates in pFus-faces/FFA-1, but not mFus-faces/FFA-2, and was recently proposed to send visual signals to dorsal retinotopic areas involved in attention and face-processing regions, as well as to re-route neural signals to allow non-serial processing among face-processing regions to expedite neural communication and protect against damage to the face processing system (Weiner et al., 2016). The vertical portion of the posterior arcuate fasciculus (pAF), on the other hand, terminates at the anterolateral tip of the mid-fusiform sulcus, which is the location of mFus-faces/FFA-2. Thus, a testable hypothesis in future studies is that the portion of the VOF terminating in pFus-faces/FFA-1 is involved in face processing and attention, while the pAF integrates semantic information and information about faces within mFus-faces/FFA-2. The integrated information would then be sent to the anterior temporal lobe through the longitudinal connections identified previously and discussed in the previous section (Gomez et al., 2015). Future studies can also integrate gyral crowns into this wiring diagram of face-selective regions, as well as shorter white matter fibers such as the recently identified fusilum (Catani, 2022) to come closer to building a mechanistic understanding among the complex relationship of cortical folding, white matter organization, face-selective regions on the FG, and face processing ability. Altogether, the present findings provide a new anatomical feature to incorporate into theories of how the human brain processes faces, as well as revealing a new neuroanatomical target in clinical populations who have difficulties processing faces.

## Acknowledgments

This research was supported by NSF CAREER Award 2042251 (Weiner), NIH Medical Scientist Training Program Grant T32 GM140935 (Willbrand), NIH T32 NS047987 (Kelly), as well as a Key Program of National Natural Science Foundation of China 62236001 (Zhen) and a General Program of National Natural Science Foundation of China 31771251 (Zhen). Data collection at Dartmouth was supported by a Rockefeller Foundation award. We thank Guo Jiahui and Hua Yang for their work with recruitment and data collection for the original project and the participants for participating.

---

**Extended Data Figure 1.**
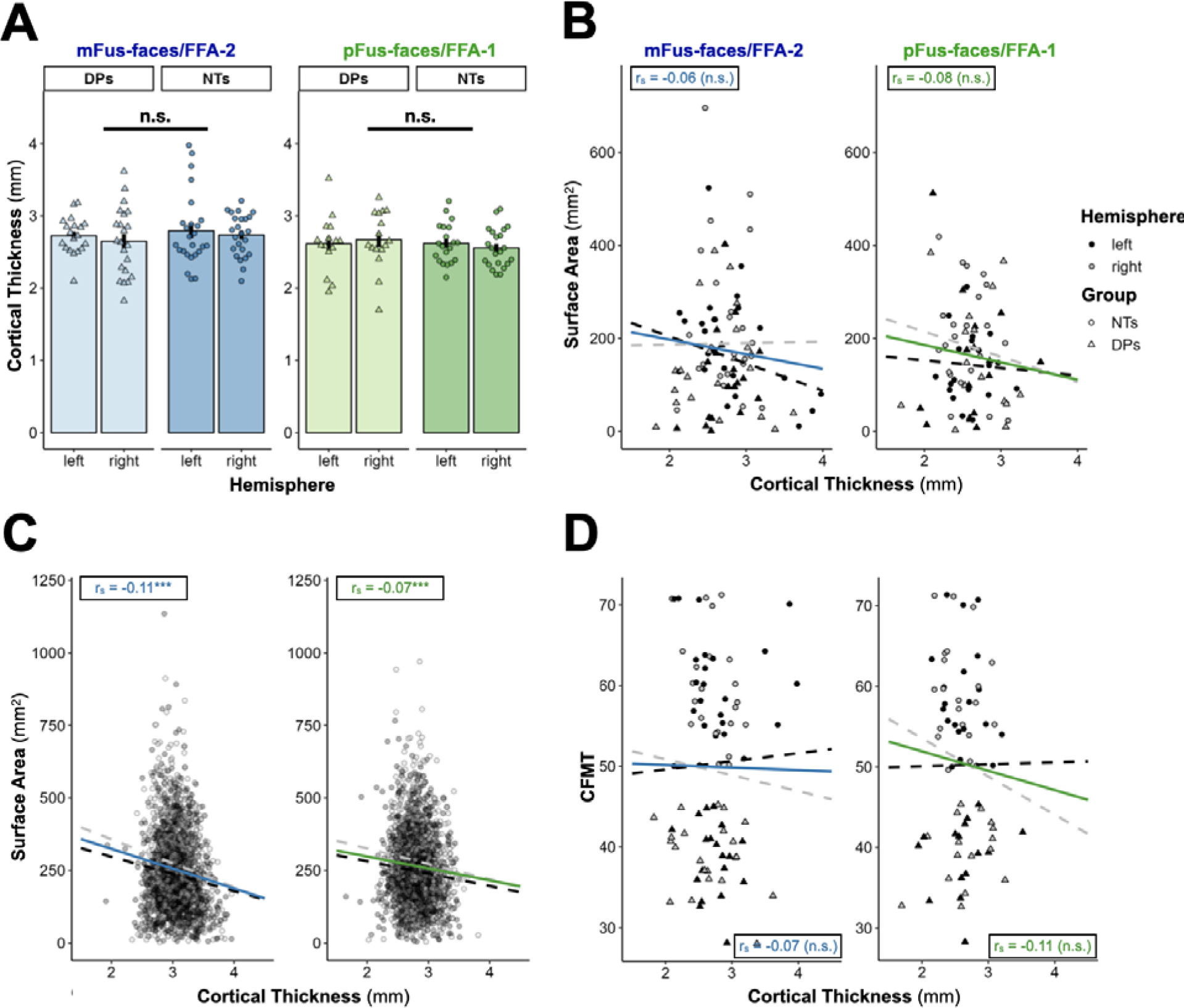
Cortical thickness of mFus and pFus does not differ between DPs and NTs and is not correlated with region size. **A.** Bar plots and error bars (mean ± se) visualizing region mean cortical thickness (in mm) as a function of hemisphere (x-axis) and group (left: DPs; right: NTs) for both mFus (left plot) and pFus (right plot). Individual dots represent individual values for each participant (DPs = triangles, NTs = circles). The line and asterisks visualize the statistical significance of the group difference for gyral crown height across hemispheres. **B.** Scatterplot of region means cortical thickness (in mm; x-axis) and surface area (in mm^2^; y-axis) for mFus (left) and pFus (right). The solid, colored line corresponds to the best-fit line across hemispheres alongside the corresponding correlation coefficient (r_s_) and p*-*values (asterisks). The best-fit lines for the left and right hemispheres are also shown as dashed lines colored according to the key. Individual dots represent individual values for each participant which are colored by hemisphere (left = black, right = white; see key) and shaped according to the group of that individual (DPs = triangles, NTs = circles; see key). **C.** Same format as B, but for the HCP sample (N = 1053) (Chen et al., 2023). **D.** Same format as B, but for the Cambridge Face Memory Test (CFMT; y-axis) performance. (*** p < .001; n.s., p > .05).

## References

Abbasi N, Duncan J, Rajimehr R (2020) Genetic influence is linked to cortical morphology in category-selective areas of visual cortex. Nat Commun 11:709 Available at: https://europepmc.org/article/pmc/pmc7002610.

Arcaro MJ, Mautz T, Berezovskii VK, Livingstone MS (2020) Anatomical correlates of face patches in macaque inferotemporal cortex. Proc Natl Acad Sci U S A 117:32667–32678 Available at: 10.1073/pnas.2018780117.

Auzias G, Lefèvre J, Le Troter A, Fischer C, Perrot M, Régis J, Coulon O (2013) Model-driven harmonic parameterization of the cortical surface: HIP-HOP. IEEE Trans Med Imaging 32:873–887 Available at: 10.1109/TMI.2013.2241651.

Avidan G, Behrmann M (2009) Functional MRI reveals compromised neural integrity of the face processing network in congenital prosopagnosia. Curr Biol 19:1146–1150 Available at: 10.1016/j.cub.2009.04.060.

Avidan G, Hasson U, Malach R, Behrmann M (2005) Detailed exploration of face-related processing in congenital prosopagnosia: 2. Functional neuroimaging findings. J Cogn Neurosci 17:1150–1167 Available at: 10.1162/0898929054475145.

Avidan G, Tanzer M, Hadj-Bouziane F, Liu N, Ungerleider LG, Behrmann M (2014) Selective dissociation between core and extended regions of the face processing network in congenital prosopagnosia. Cereb Cortex 24:1565–1578 Available at: 10.1093/cercor/bht007.

Barch DM et al. (2013) Function in the human connectome: task-fMRI and individual differences in behavior. Neuroimage 80:169–189 Available at: 10.1016/j.neuroimage.2013.05.033.

Behrmann M, Scherf KS, Avidan G (2016) Neural mechanisms of face perception, their emergence over development, and their breakdown. Wiley Interdiscip Rev Cogn Sci 7:247– 263 Available at: 10.1002/wcs.1388.

Bi T, Chen J, Zhou T, He Y, Fang F (2014) Function and structure of human left fusiform cortex are closely associated with perceptual learning of faces. Curr Biol 24:222–227 Available at: 10.1016/j.cub.2013.12.028.

Bouhali F, Dubois J, Hoeft F, Weiner KS (2024) Unique longitudinal contributions of sulcal interruptions to reading acquisition in children. eLife Available at: https://elifesciences.org/reviewed-preprints/103007 [Accessed January 29, 2025].

Cachia A, Roell M, Mangin J-F, Sun ZY, Jobert A, Braga L, Houde O, Dehaene S, Borst G (2018) How interindividual differences in brain anatomy shape reading accuracy. Brain Struct Funct 223:701–712 Available at: 10.1007/s00429-017-1516-x.

Catani M (2022) The connectional anatomy of the temporal lobe. Handb Clin Neurol 187:3–16 Available at: 10.1016/B978-0-12-823493-8.00001-8.

Chen X, Liu X, Parker BJ, Zhen Z, Weiner KS (2023) Functionally and structurally distinct fusiform face area(s) in over 1000 participants. Neuroimage 265:119765 Available at: 10.1016/j.neuroimage.2022.119765.

Cottaar M, Bastiani M, Boddu N, Glasser MF, Haber S, van Essen DC, Sotiropoulos SN, Jbabdi S (2021) Modelling white matter in gyral blades as a continuous vector field. Neuroimage 227:117693 Available at: 10.1016/j.neuroimage.2020.117693.

Çukur T, Huth AG, Nishimoto S, Gallant JL (2013) Functional subdomains within human FFA. J Neurosci 33:16748–16766 Available at: 10.1523/JNEUROSCI.1259-13.2013.

Dale AM, Fischl B, Sereno MI (1999) Cortical surface-based analysis. I. Segmentation and surface reconstruction. Neuroimage 9:179–194 Available at: 10.1006/nimg.1998.0395.

Davidenko N, Remus DA, Grill-Spector K (2012) Face-likeness and image variability drive responses in human face-selective ventral regions. Hum Brain Mapp 33:2334–2349 Available at: 10.1002/hbm.21367.

Duchaine B, Germine L, Nakayama K (2007a) Family resemblance: ten family members with prosopagnosia and within-class object agnosia. Cogn Neuropsychol 24:419–430 Available at: 10.1080/02643290701380491.

Duchaine B, Nakayama K (2005) Dissociations of face and object recognition in developmental prosopagnosia. J Cogn Neurosci 17:249–261 Available at: 10.1162/0898929053124857.

Duchaine B, Nakayama K (2006) The Cambridge Face Memory Test: results for neurologically intact individuals and an investigation of its validity using inverted face stimuli and prosopagnosic participants. Neuropsychologia 44:576–585 Available at: 10.1016/j.neuropsychologia.2005.07.001.

Duchaine B, Yovel G (2015) A Revised Neural Framework for Face Processing. Annu Rev Vis Sci 1:393–416 Available at: 10.1146/annurev-vision-082114-035518.

Duchaine B, Yovel G, Nakayama K (2007b) No global processing deficit in the Navon task in 14 developmental prosopagnosics. Soc Cogn Affect Neurosci 2:104–113 Available at: 10.1093/scan/nsm003.

Elbich DB, Scherf S (2017) Beyond the FFA: Brain-behavior correspondences in face recognition abilities. Neuroimage 147:409–422 Available at: 10.1016/j.neuroimage.2016.12.042.

Engell AD, McCarthy G (2013) Probabilistic atlases for face and biological motion perception: an analysis of their reliability and overlap. Neuroimage 74:140–151 Available at: 10.1016/j.neuroimage.2013.02.025.

Finzi D, Gomez J, Nordt M, Rezai AA, Poltoratski S, Grill-Spector K (2021) Differential spatial computations in ventral and lateral face-selective regions are scaffolded by structural connections. Nat Commun 12:2278 Available at: 10.1038/s41467-021-22524-2.

Fischl B, Dale AM (2000) Measuring the thickness of the human cerebral cortex from magnetic resonance images. Proc Natl Acad Sci U S A 97:11050–11055 Available at: 10.1073/pnas.200033797.

Fischl B, Sereno MI, Dale AM (1999a) Cortical surface-based analysis. II: Inflation, flattening, and a surface-based coordinate system. Neuroimage 9:195–207 Available at: 10.1006/nimg.1998.0396.

Fischl B, Sereno MI, Tootell RB, Dale AM (1999b) High-resolution intersubject averaging and a coordinate system for the cortical surface. Hum Brain Mapp 8:272–284 Available at: https://onlinelibrary.wiley.com/doi/10.1002/(SICI)1097-0193(1999)8:4%3C272::AID-HBM10%3E3.0.CO;2-4.

Furl N, Garrido L, Dolan RJ, Driver J, Duchaine B (2011) Fusiform gyrus face selectivity relates to individual differences in facial recognition ability. J Cogn Neurosci 23:1723–1740 Available at: 10.1162/jocn.2010.21545.

Garrido L, Furl N, Draganski B, Weiskopf N, Stevens J, Tan GC-Y, Driver J, Dolan RJ, Duchaine B (2009) Voxel-based morphometry reveals reduced grey matter volume in the temporal cortex of developmental prosopagnosics. Brain 132:3443–3455 Available at: 10.1093/brain/awp271.

Glasser MF, Sotiropoulos SN, Wilson JA, Coalson TS, Fischl B, Andersson JL, Xu J, Jbabdi S, Webster M, Polimeni JR, Van Essen DC, Jenkinson M, WU-Minn HCP Consortium (2013) The minimal preprocessing pipelines for the Human Connectome Project. Neuroimage 80:105–124 Available at: 10.1016/j.neuroimage.2013.04.127.

Golarai G, Ghahremani DG, Whitfield-Gabrieli S, Reiss A, Eberhardt JL, Gabrieli JDE, Grill-Spector K (2007) Differential development of high-level visual cortex correlates with category-specific recognition memory. Nat Neurosci 10:512–522 Available at: 10.1038/nn1865.

Golarai G, Hong S, Haas BW, Galaburda AM, Mills DL, Bellugi U, Grill-Spector K, Reiss AL (2010) The fusiform face area is enlarged in Williams syndrome. J Neurosci 30:6700–6712 Available at: 10.1523/JNEUROSCI.4268-09.2010.

Gomez J, Barnett MA, Natu V, Mezer A, Palomero-Gallagher N, Weiner KS, Amunts K, Zilles K, Grill-Spector K (2017) Microstructural proliferation in human cortex is coupled with the development of face processing. Science 355:68–71 Available at: 10.1126/science.aag0311.

Gomez J, Natu V, Jeska B, Barnett M, Grill-Spector K (2018) Development differentially sculpts receptive fields across early and high-level human visual cortex. Nat Commun 9:788 Available at: 10.1038/s41467-018-03166-3.

Gomez J, Pestilli F, Witthoft N, Golarai G, Liberman A, Poltoratski S, Yoon J, Grill-Spector K (2015) Functionally defined white matter reveals segregated pathways in human ventral temporal cortex associated with category-specific processing. Neuron 85:216–227 Available at: 10.1016/j.neuron.2014.12.027.

Jiahui G, Yang H, Duchaine B (2018) Developmental prosopagnosics have widespread selectivity reductions across category-selective visual cortex. Proc Natl Acad Sci U S A 115:E6418–E6427 Available at: 10.1073/pnas.1802246115.

Jonas J, Brissart H, Hossu G, Colnat-Coulbois S, Vignal J-P, Rossion B, Maillard L (2018) A face identity hallucination (palinopsia) generated by intracerebral stimulation of the face-selective right lateral fusiform cortex. Cortex 99:296–310 Available at: 10.1016/j.cortex.2017.11.022.

Jonas J, Rossion B (2021) Intracerebral electrical stimulation to understand the neural basis of human face identity recognition. Eur J Neurosci Available at: 10.1111/ejn.15235.

Julian JB, Fedorenko E, Webster J, Kanwisher N (2012) An algorithmic method for functionally defining regions of interest in the ventral visual pathway. Neuroimage 60:2357–2364 Available at: 10.1016/j.neuroimage.2012.02.055.

Kanwisher N (2010) Functional specificity in the human brain: a window into the functional architecture of the mind. Proc Natl Acad Sci U S A 107:11163–11170 Available at: 10.1073/pnas.1005062107.

Kanwisher N, McDermott J, Chun MM (1997) The fusiform face area: a module in human extrastriate cortex specialized for face perception. J Neurosci 17:4302–4311 Available at: 10.1523/JNEUROSCI.17-11-04302.1997.

Kay KN, Weiner KS, Grill-Spector K (2015) Attention reduces spatial uncertainty in human ventral temporal cortex. Curr Biol 25:595–600 Available at: 10.1016/j.cub.2014.12.050.

Kietzmann TC, Swisher JD, König P, Tong F (2012) Prevalence of selectivity for mirror-symmetric views of faces in the ventral and dorsal visual pathways. J Neurosci 32:11763– 11772 Available at: 10.1523/JNEUROSCI.0126-12.2012.

Kriegeskorte N, Simmons WK, Bellgowan PSF, Baker CI (2009) Circular analysis in systems neuroscience: the dangers of double dipping. Nat Neurosci 12:535–540 Available at: 10.1038/nn.2303.

McGugin RW, Newton AT, Gore JC, Gauthier I (2014) Robust expertise effects in right FFA. Neuropsychologia 63:135–144 Available at: 10.1016/j.neuropsychologia.2014.08.029.

McGugin RW, Newton AT, Tamber-Rosenau B, Tomarken A, Gauthier I (2020) Thickness of Deep Layers in the Fusiform Face Area Predicts Face Recognition. J Cogn Neurosci 32:1316–1329 Available at: 10.1162/jocn_a_01551.

McGugin RW, Van Gulick AE, Gauthier I (2016) Cortical Thickness in Fusiform Face Area Predicts Face and Object Recognition Performance. J Cogn Neurosci 28:282–294 Available at: 10.1162/jocn_a_00891.

McGugin RW, Van Gulick AE, Tamber-Rosenau BJ, Ross DA, Gauthier I (2015) Expertise Effects in Face-Selective Areas are Robust to Clutter and Diverted Attention, but not to Competition. Cereb Cortex 25:2610–2622 Available at: 10.1093/cercor/bhu060.

Miller JA, Voorhies WI, Lurie DJ, D’Esposito M, Weiner KS (2021) Overlooked Tertiary Sulci Serve as a Meso-Scale Link between Microstructural and Functional Properties of Human Lateral Prefrontal Cortex. J Neurosci 41:2229–2244 Available at: https://www.jneurosci.org/content/41/10/2229 [Accessed July 19, 2021].

Natu VS, Arcaro MJ, Barnett MA, Gomez J, Livingstone M, Grill-Spector K, Weiner KS (2021) Sulcal Depth in the Medial Ventral Temporal Cortex Predicts the Location of a Place-Selective Region in Macaques, Children, and Adults. Cereb Cortex 31:48–61 Available at: 10.1093/cercor/bhaa203.

Natu VS, Barnett MA, Hartley J, Gomez J, Stigliani A, Grill-Spector K (2016) Development of Neural Sensitivity to Face Identity Correlates with Perceptual Discriminability. J Neurosci 36:10893–10907 Available at: 10.1523/JNEUROSCI.1886-16.2016.

Natu VS, Gomez J, Barnett M, Jeska B, Kirilina E, Jaeger C, Zhen Z, Cox S, Weiner KS, Weiskopf N, Grill-Spector K (2019) Apparent thinning of human visual cortex during childhood is associated with myelination. Proc Natl Acad Sci U S A 116:20750–20759 Available at: 10.1073/pnas.1904931116.

Nichols TE, Poline J-B (2009) Commentary on Vul et al.’s (2009) “Puzzlingly High Correlations in fMRI Studies of Emotion, Personality, and Social Cognition.” Perspect Psychol Sci 4:291–293 Available at: 10.1111/j.1745-6924.2009.01126.x.

Nordt M, Gomez J, Natu VS, Rezai AA, Finzi D, Kular H, Grill-Spector K (2021) Cortical recycling in high-level visual cortex during childhood development. Nat Hum Behav 5:1686– 1697 Available at: 10.1038/s41562-021-01141-5.

Norman-Haignere S, Kanwisher N, McDermott JH (2013) Cortical pitch regions in humans respond primarily to resolved harmonics and are located in specific tonotopic regions of anterior auditory cortex. J Neurosci 33:19451–19469 Available at: 10.1523/JNEUROSCI.2880-13.2013.

Norman-Haignere SV, Albouy P, Caclin A, McDermott JH, Kanwisher NG, Tillmann B (2016) Pitch-Responsive Cortical Regions in Congenital Amusia. J Neurosci 36:2986–2994 Available at: 10.1523/JNEUROSCI.2705-15.2016.

Parker BJ, Voorhies WI, Jiahui G, Miller JA, Willbrand E, Hallock T, Furl N, Garrido L, Duchaine B, Weiner KS (2023) Hominoid-specific sulcal variability is related to face perception ability. Brain Struct Funct 228:677–685 Available at: 10.1007/s00429-023-02611-4.

Parvizi J, Jacques C, Foster BL, Withoft N, Rangarajan V, Weiner KS, Grill-Spector K (2012) Electrical Stimulation of Human Fusiform Face-Selective Regions Distorts Face Perception. Journal of Neuroscience 32:14915–14920 Available at: 10.1523/jneurosci.2609-12.2012.

Pinsk MA, Arcaro M, Weiner KS, Kalkus JF, Inati SJ, Gross CG, Kastner S (2009) Neural representations of faces and body parts in macaque and human cortex: a comparative FMRI study. J Neurophysiol 101:2581–2600 Available at: 10.1152/jn.91198.2008.

Poldrack RA, Mumford JA (2009) Independence in ROI analysis: where is the voodoo? Soc Cogn Affect Neurosci 4:208–213 Available at: 10.1093/scan/nsp011.

Rangarajan V, Hermes D, Foster BL, Weiner KS, Jacques C, Grill-Spector K, Parvizi J (2014) Electrical stimulation of the left and right human fusiform gyrus causes different effects in conscious face perception. J Neurosci 34:12828–12836 Available at: 10.1523/JNEUROSCI.0527-14.2014.

Rosenke M, Davidenko N, Grill-Spector K, Weiner KS (2020) Combined Neural Tuning in Human Ventral Temporal Cortex Resolves the Perceptual Ambiguity of Morphed 2D Images. Cereb Cortex 30:4882–4898 Available at: 10.1093/cercor/bhaa081.

Rosenke M, van Hoof R, van den Hurk J, Grill-Spector K, Goebel R (2021) A Probabilistic Functional Atlas of Human Occipito-Temporal Visual Cortex. Cereb Cortex 31:603–619 Available at: 10.1093/cercor/bhaa246.

Schalk G, Kapeller C, Guger C, Ogawa H, Hiroshima S, Lafer-Sousa R, Saygin ZM, Kamada K, Kanwisher N (2017) Facephenes and rainbows: Causal evidence for functional and anatomical specificity of face and color processing in the human brain. Proc Natl Acad Sci U S A 114:12285–12290 Available at: 10.1073/pnas.1713447114.

Scherf KS, Elbich DB, Motta-Mena NV (2017) Investigating the Influence of Biological Sex on the Behavioral and Neural Basis of Face Recognition. eNeuro 4 Available at: 10.1523/ENEURO.0104-17.2017.

Schrouff J, Raccah O, Baek S, Rangarajan V, Salehi S, Mourão-Miranda J, Helili Z, Daitch AL, Parvizi J (2020) Fast temporal dynamics and causal relevance of face processing in the human temporal cortex. Nat Commun 11:656 Available at: 10.1038/s41467-020-14432-8.

Stigliani A, Jeska B, Grill-Spector K (2019) Differential sustained and transient temporal processing across visual streams. PLoS Comput Biol 15:e1007011 Available at: 10.1371/journal.pcbi.1007011.

Stigliani A, Weiner KS, Grill-Spector K (2015) Temporal Processing Capacity in High-Level Visual Cortex Is Domain Specific. J Neurosci 35:12412–12424 Available at: 10.1523/JNEUROSCI.4822-14.2015.

Susilo T, Duchaine B (2013) Advances in developmental prosopagnosia research. Curr Opin Neurobiol 23:423–429 Available at: 10.1016/j.conb.2012.12.011.

Tian X, Wang R, Zhao Y, Zhen Z, Song Y, Liu J (2020) Multi-item discriminability pattern to faces in developmental prosopagnosia reveals distinct mechanisms of face processing. Cereb Cortex 30:2986–2996 Available at: https://pubmed.ncbi.nlm.nih.gov/31813985/ [Accessed January 29, 2025].

Van Essen DC et al. (2012) The Human Connectome Project: a data acquisition perspective. Neuroimage 62:2222–2231 Available at: 10.1016/j.neuroimage.2012.02.018.

Van Essen DC, Jbabdi S, Sotiropoulos SN, Chen C, Dikranian K, Coalson T, Harwell J, Behrens TEJ, Glasser MF (2014) Chapter 16 - Mapping Connections in Humans and Non-Human Primates: Aspirations and Challenges for Diffusion Imaging. In: Diffusion MRI (Second Edition) (Johansen-Berg H, Behrens TEJ, eds), pp 337–358. San Diego: Academic Press. Available at: https://www.sciencedirect.com/science/article/pii/B9780123964601000160.

Vul E, Harris C, Winkielman P, Pashler H (2009) Puzzlingly High Correlations in fMRI Studies of Emotion, Personality, and Social Cognition. Perspect Psychol Sci 4:274–290 Available at: 10.1111/j.1745-6924.2009.01125.x.

Weiner KS (2019) The Mid-Fusiform Sulcus (sulcus sagittalis gyri fusiformis). Anat Rec 302:1491–1503 Available at: https://onlinelibrary.wiley.com/doi/10.1002/ar.24041.

Weiner KS, Barnett MA, Lorenz S, Caspers J, Stigliani A, Amunts K, Zilles K, Fischl B, Grill-Spector K (2017a) The Cytoarchitecture of Domain-specific Regions in Human High-level Visual Cortex. Cereb Cortex 27:146–161 Available at: 10.1093/cercor/bhw361.

Weiner KS, Golarai G, Caspers J, Chuapoco MR, Mohlberg H, Zilles K, Amunts K, Grill-Spector K (2014) The mid-fusiform sulcus: a landmark identifying both cytoarchitectonic and functional divisions of human ventral temporal cortex. Neuroimage 84:453–465 Available at: 10.1016/j.neuroimage.2013.08.068.

Weiner KS, Grill-Spector K (2010) Sparsely-distributed organization of face and limb activations in human ventral temporal cortex. Neuroimage 52:1559–1573 Available at: 10.1016/j.neuroimage.2010.04.262.

Weiner KS, Jonas J, Gomez J, Maillard L, Brissart H, Hossu G, Jacques C, Loftus D, Colnat-Coulbois S, Stigliani A, Barnett MA, Grill-Spector K, Rossion B (2016) The Face-Processing Network Is Resilient to Focal Resection of Human Visual Cortex. J Neurosci 36:8425–8440 Available at: 10.1523/JNEUROSCI.4509-15.2016.

Weiner KS, Sayres R, Vinberg J, Grill-Spector K (2010) fMRI-adaptation and category selectivity in human ventral temporal cortex: regional differences across time scales. J Neurophysiol 103:3349–3365 Available at: 10.1152/jn.01108.2009.

Weiner KS, Willbrand EH (2023) Is there an Association between Tuber Involvement of the Fusiform Face Area in Autism Diagnosis? Ann Neurol 93:1218–1220 Available at: 10.1002/ana.26632.

Weiner KS, Yeatman JD, Wandell BA (2017b) The posterior arcuate fasciculus and the vertical occipital fasciculus. Cortex 97:274–276 Available at: 10.1016/j.cortex.2016.03.012.

Wilmer JB, Germine L, Chabris CF, Chatterjee G, Gerbasi M, Nakayama K (2012) Capturing specific abilities as a window into human individuality: the example of face recognition. Cogn Neuropsychol 29:360–392 Available at: 10.1080/02643294.2012.753433.

Wilmer JB, Germine L, Chabris CF, Chatterjee G, Williams M, Loken E, Nakayama K, Duchaine B (2010) Human face recognition ability is specific and highly heritable. Proc Natl Acad Sci U S A 107:5238–5241 Available at: 10.1073/pnas.0913053107.

Witthoft N, Poltoratski S, Nguyen M, Golarai G, Liberman A, LaRocque KF, Smith ME, Grill-Spector K (2016) Reduced spatial integration in the ventral visual cortex underlies face recognition deficits in developmental prosopagnosia. bioRxiv:051102 Available at: https://www.biorxiv.org/content/10.1101/051102v1.full [Accessed January 31, 2023].

Yeatman JD, Weiner KS, Pestilli F, Rokem A, Mezer A, Wandell BA (2014) The vertical occipital fasciculus: a century of controversy resolved by in vivo measurements. Proc Natl Acad Sci U S A 111:E5214–E5223 Available at: 10.1073/pnas.1418503111.

Zebrowitz L, Ward N, Boshyan J, Gutchess A, Hadjikhani N (2016) Dedifferentiated face processing in older adults is linked to lower resting state metabolic activity in fusiform face area. Brain Res 1644:22–31 Available at: 10.1016/j.brainres.2016.05.007.

Zhang J, Liu J, Xu Y (2015) Neural decoding reveals impaired face configural processing in the right fusiform face area of individuals with developmental prosopagnosia. J Neurosci 35:1539–1548 Available at: 10.1523/JNEUROSCI.2646-14.2015.

Zhang S, Chavoshnejad P, Li X, Guo L, Jiang X, Han J, Wang L, Li G, Wang X, Liu T, Razavi MJ, Zhang S, Zhang T (2022) Gyral peaks: Novel gyral landmarks in developing macaque brains. Hum Brain Mapp 43:4540–4555 Available at: 10.1002/hbm.25971.

Zhang S, Zhang T, Cao G, Zhou J, He Z, Li X, Ren Y, Jiang X, Guo L, Han J, Liu T (2023) Species-Shared and -Unique Gyral Peaks on Human and Macaque Brains. bioRxiv Available at: 10.1101/2023.07.26.550760.

Zhao Y, Li J, Liu X, Song Y, Wang R, Yang Z, Liu J (2016) Altered spontaneous neural activity in the occipital face area reflects behavioral deficits in developmental prosopagnosia. Neuropsychologia 89:344–355 Available at: 10.1016/j.neuropsychologia.2016.05.027 [Accessed January 29, 2025].

Zhao Y, Zhen Z, Liu X, Song Y, Liu J (2018) The neural network for face recognition: Insights from an fMRI study on developmental prosopagnosia. Neuroimage 169:151–161 Available at: 10.1016/j.neuroimage.2017.12.023 [Accessed January 29, 2025].

Zhen Z, Yang Z, Huang L, Kong X-Z, Wang X, Dang X, Huang Y, Song Y, Liu J (2015) Quantifying interindividual variability and asymmetry of face-selective regions: a probabilistic functional atlas. Neuroimage 113:13–25 Available at: 10.1016/j.neuroimage.2015.03.010.

